# The IMPDH inhibitor merimepodib suppresses SARS-CoV-2 replication *in vitro*

**DOI:** 10.1101/2020.04.07.028589

**Authors:** Natalya Bukreyeva, Emily K. Mantlo, Rachel A. Sattler, Cheng Huang, Slobodan Paessler, Jerry Zeldis

## Abstract

The ongoing COVID-19 pandemic continues to pose a major public health burden around the world. The novel coronavirus, severe acute respiratory syndrome coronavirus 2 (SARS-CoV-2), has infected over one million people worldwide as of April, 2020, and has led to the deaths of nearly 300,000 people. No approved vaccines or treatments in the USA currently exist for COVID-19, so there is an urgent need to develop effective countermeasures. The IMPDH inhibitor merimepodib (MMPD) is an investigational antiviral drug that acts as a noncompetitive inhibitor of IMPDH. It has been demonstrated to suppress replication of a variety of emerging RNA viruses. We report here that MMPD suppresses SARS-CoV-2 replication in vitro. After overnight pretreatment of Vero cells with 10 μM of MMPD, viral titers were reduced by 4 logs of magnitude, while pretreatment for 4 hours resulted in a 3-log drop. The effect is dose-dependent, and concentrations as low as 3.3 μM significantly reduced viral titers when the cells were pretreated prior to infection. The results of this study provide evidence that MMPD may be a viable treatment option for COVID-19.

## Introduction

In December of 2019, a novel coronavirus emerged out of Wuhan, China, causing respiratory illness and occasional severe pneumonia. The virus spread rapidly around the world, and in March of 2019 the disease was declared a pandemic by the World Health Organization. At the time of writing, 1.3 million people around the world have been infected and more than 70,000 people have died from the disease, named coronavirus disease 2019 (COVID-19). In addition to the public health crisis, the economic impact from the pandemic continues to worsen. Due to the novelty of the virus, no approved vaccines or treatments are available. There is therefore an urgent need for the development of new countermeasures.

Drugs with history of being tested in human patients or used for treatment of other conditions offer the most expedient option, and several such drugs are currently being tested for efficacy against SARS-CoV-2, including many broad-spectrum antivirals. One such antiviral, merimepodib (MMPD), has already being tested against hepatitis C in patients as well as against many emerging RNA viruses in cell culture, including Zika, Ebola, Lassa, Junin, and chikungunya viruses (1). MMPD noncompetitively inhibits inosine-5’-monophosphate dehydrogenase (IMPDH), an enzyme responsible for *de novo* synthesis of guanosine nucleotides (2, 3). *In vitro*, inhibition can be reversed by addition of exogenous guanosine (4). Early work suggests that IMPDH may directly interact with SARS-CoV-2 nsp13, perhaps indicating that drugs targeting IMPDH such as MMPD might have an impact on viral replication (5). This drug, while still investigational, is considered to be safe in humans. More than 300 patients have received the drug in Phase I and Phase II clinical trials. Many were treated for six months. In this study, we aimed to examine whether MMPD may be active in reducing SARS-CoV-2 replication.

## Materials & Methods

### Cells and Viruses

Vero cells (CCL-81, ATCC) were maintained in Dulbecco’s modified eagle medium (Hyclone) supplemented with 10% FBS (Gibco) and 1% penicillin and streptomycin solution (Hyclone). SARS-CoV-2 USA-WA1/2020 was provided by the World Reference Center for Emerging Viruses and Arboviruses (WRCEVA) and was originally obtained from the Centers for Disease Control and Prevention (CDC). The virus was subsequently titrated and passaged once more on Vero cells, grown in DMEM supplemented with 5% FBS and 1% penicillin and streptomycin solution. All experiments were conducted at the University of Texas Medical Branch (UTMB) approved biosafety level-three (BSL-3) laboratories and all personnel undergo routine medical surveillance.

### Compound pretreatment and infection

Vero cells were pretreated, 1 mL per well, with DMEM media containing the indicated concentrations of either merimepodib or T-705 and incubated at 37°C for either 4 hours or overnight. The media was then removed and cells were inoculated with SARS-CoV-2 at an MOI of 0.05. Viral inoculum was premixed with compounds (0.1 mL/well). After 1 hour of incubation at 37°C, wells were washed 3 times with DMEM media and 1 mL of media containing the indicated compound doses was added back to each well. Cells were incubated at 37°C and samples collected at 0, 16, and 24 hours post-infection. Collected timepoint media was replaced with an equal amount of media containing compound. The SARS-CoV-2 titer was determined for each sample collected at 0, 16, and 24 hours post-infection via TCID_50_ in Vero cells. Briefly, collected supernatant was serially diluted six times (10^−6^) in DMEM media. 100 μl of each dilution was added in quadruplicates to Vero cells grown in a 96-well plate. Cells were incubated 4 days at 37°C. Cells were then fixed with 10% formalin for 30 minutes and stained for 5 minutes with crystal violet. Biological triplicates were harvested at each timepoint and statistical significance was determined via one-tailed t-test.

## Results

Previous growth kinetics studies using an MOI of 0.01 in Vero cells have indicated that viral titers peak at around 24 hours post-infection and plateau through 48 hours post-infection (6). At both 16 and 24 hours post-infection, during the exponential phase of the viral growth curve, 10 μM MMPD reduced viral titers by around 3 logs when cells were pretreated overnight (p-value = <0.0001). 4 hours of pretreatment with MMPD resulted in a nearly 2.5-log decrease in viral titers (p-value = 0.0004). The antiviral effect is dose-dependent, with lower doses of 5 and 3.3 μM of MMPD reducing viral titers by a more modest, but still significant degree. At 24 hours post-infection, 5 μM MMPD applied 4 hours before infection reduced viral titers by 1.5 logs (p-value = 0.0004), while 3.3 μM added 4 hours before infection reduced viral titers by a little over 1 log (p-value = 0.001).

We also tested T-705, another broad-spectrum antiviral that acts as a nucleoside analog. Both overnight and 4-hour pretreatment with high doses (33-100 μM) of T-705 failed to inhibit SARS-CoV-2 replication. Overall, our results indicate that therapeutic concentrations of MMPD, but not T-705, is effective in reducing SARS-COV-2 coronavirus titers.

## Discussion

Our results show that MMPD can inhibit SARS-CoV-2 replication at low concentrations. This is likely due its inhibition of IMPDH, leading to a depletion of guanosine for use by the viral polymerase during replication. By contrast, T-705 has been reported to weakly inhibit IMPDH, instead acting as a nucleotide analogue and interacting more specifically with certain viral polymerases (7). Further work is needed to characterize the full mechanism behind MMPD inhibition of SARS-CoV-2 as well as its efficacy in animal models of corona virus infections.

In COVID-19 infection it is essential to minimize the spread of the viral infection to the lower respiratory tract. We chose to test MMPD pretreated uninfected cells in order to see if viral spread can be limited. This approach potentially allows us to model the use of this drug for prophylaxis. This is important since a large proportion of exposed patients can experience rapid expansion of their viral burden while being asymptomatic. A drug that thwarts this viral expansion will allow the immune system to eliminate the nascent infection. Since MMPD is a host directed therapy and not a direct acting antiviral, the likelihood of emergence of resistant variants of the virus to MMPD is low. Potentially MMPD might be used in combination with direct antivirals or immunomodulators.

The concentrations tested in this study are easily clinically-achievable in human patients. 50 mg MMPD administered orally results in plasma concentrations of around 1154 ng/ml (2500 μM) shortly after administration (8). MMPD may therefore be a viable treatment option for COVID-19 that can be quickly tested and deployed.

## Figure Legends

**Figure 1:**
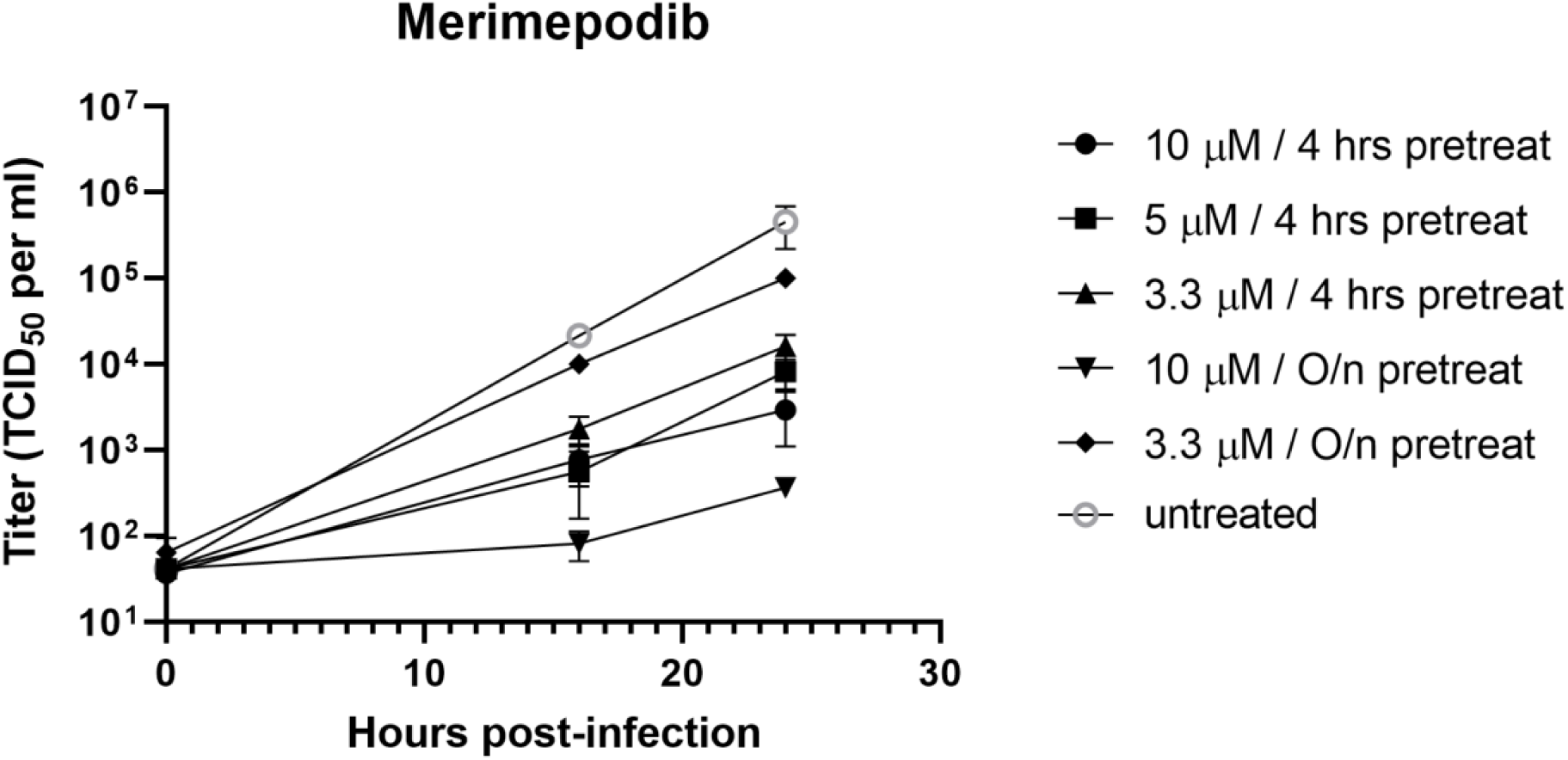
Effect of merimepodib on SARS-CoV-2 replication. Vero cells were pretreated, 1 mL per well, with the merimepodib doses listed and incubated at 37C either 4 hours or overnight. The media was then removed and cells were inoculated with 0.05 MOI of SARS-CoV-2 premixed with compounds (0.1 mL/well). After 1 hour of incubation at 37C, wells were washed 3 times with dilution media and 1 mL per well of each compound dose was added. Cells were incubated at 37C and samples collected and titrated via TCID_50_ at 0, 16, and 24 hours post-infection. The average of triplicates and standard deviation are shown.

**Figure 2:**
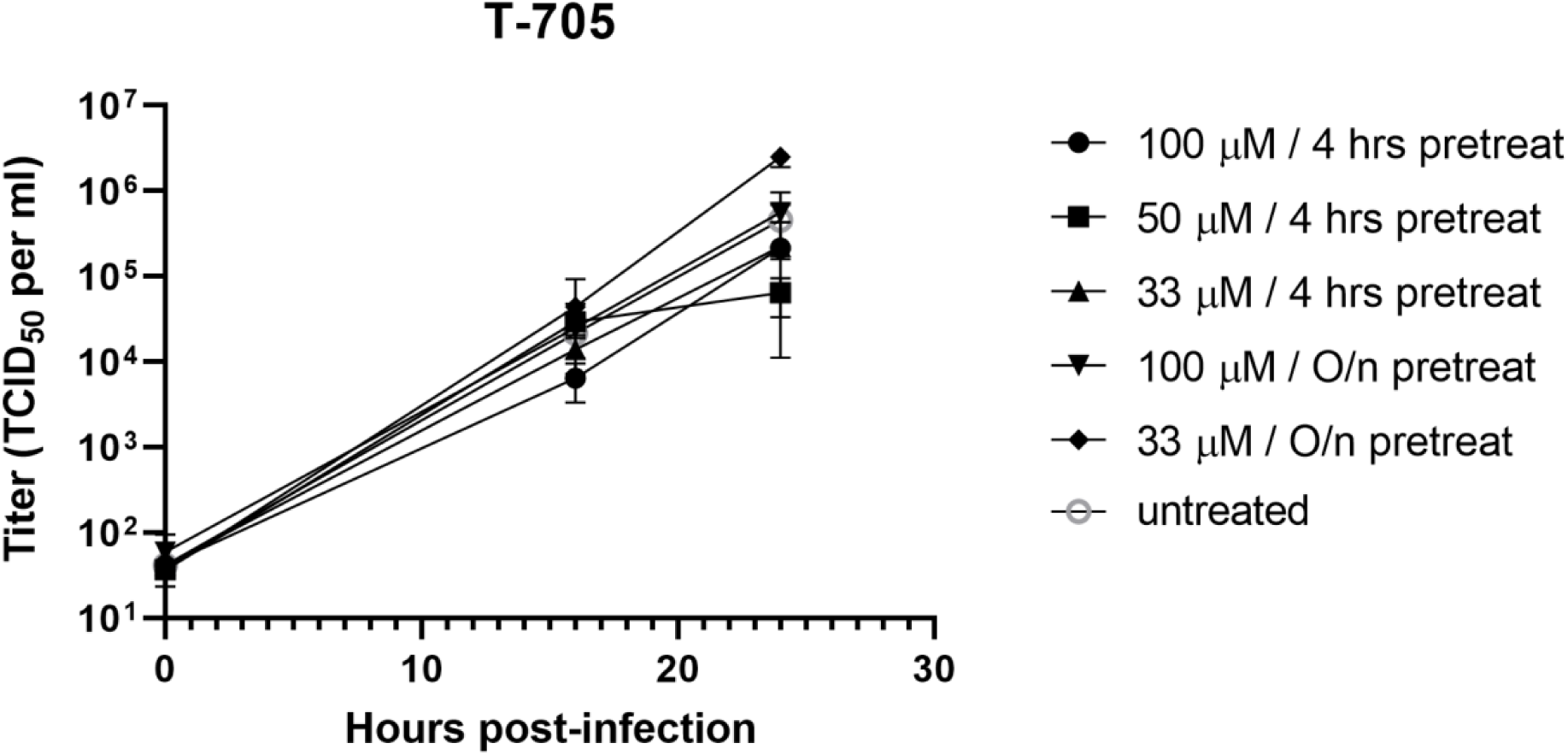
Effect of T-705 on SARS-CoV-2 replication. Vero cells were pretreated, 1 mL per well, with the T-705 doses listed and incubated at 37C either 4 hours or overnight. The media was then removed and cells were inoculated with 0.05 MOI of SARS-CoV-2 premixed with compounds (0.1 mL/well). After 1 hour of incubation at 37C, wells were washed 3 times with dilution media and 1 mL per well of each compound dose was added. Cells were incubated at 37C and samples collected and titrated via TCID_50_ at 0, 16, and 24 hours post-infection. The average of triplicates and standard deviation are shown.

